# Adjusted vascular contractility relies on integrity of progranulin pathway: Insights into mitochondrial function

**DOI:** 10.1101/2023.10.27.564485

**Authors:** Shubhnita Singh, Ariane Bruder-Nascimento, Rafael M. Costa, Juliano V. Alves, Sivakama Bharathi, Eric S. Goetzman, Thiago Bruder-Nascimento

**Affiliations:** Department of Pediatrics at UPMC Children’s Hospital of Pittsburgh, University of Pittsburgh, Pittsburgh, PA, USA; Center for Pediatrics Research in Obesity and Metabolism (CPROM) at UPMC Children’s Hospital of Pittsburgh, University of Pittsburgh, Pittsburgh, PA, USA; Endocrinology Division at UPMC Children’s Hospital of Pittsburgh, University of Pittsburgh, Pittsburgh, PA, USA; Department of Human Genetics, School of Public Health, University of Pittsburgh, Pittsburgh, USA; Vascular Medicine Institute (VMI), University of Pittsburgh, Pittsburgh, PA, USA; Genetic and Genomic Medicine Division at UPMC Children’s Hospital of Pittsburgh, University of Pittsburgh, Pittsburgh, PA, USA; Department of Pharmacology, Ribeirao Preto Medical School, University of Sao Paulo, Ribeirao Preto, SP, Brazil

**Keywords:** Progranulin, mitochondria, Reactive Oxygen Species, Vascular contractility

## Abstract

**Objective:** Cardiovascular disease (CVD) is a global health crisis and a leading cause of mortality. The intricate interplay between vascular contractility and mitochondrial function is central to CVD pathogenesis. The progranulin gene (GRN) encodes glycoprotein progranulin (PGRN), a ubiquitous molecule with known anti-inflammatory property. However, the role of PGRN in CVD remains enigmatic. In this study, we sought to dissect the significance of PGRN in the regulation vascular contractility and investigate the interface between PGRN and mitochondrial quality.

**Method:** Our investigation utilized aortae from male and female C57BL6/J wild-type (PGRN+/+) and B6(Cg)-Grntm1.1Aidi/J (PGRN−/−) mice, encompassing wire myograph assays to assess vascular contractility and primary aortic vascular smooth muscle cells (VSMCs) for mechanistic insights.

**Results:** Our results showed suppression of contractile activity in PGRN-/- VSMCs and aorta, followed by reduced α-smooth muscle actin expression. Mechanistically, PGRN deficiency impaired mitochondrial oxygen consumption rate (OCR), complex I activity, mitochondrial turnover, and mitochondrial redox signaling, while restoration of PGRN levels in aortae from PGRN-/- mice via lentivirus delivery ameliorated contractility and boosted OCR. In addition, VSMC overexpressing PGRN displayed higher mitochondrial respiration and complex I activity accompanied by cellular hypercontractility. Furthermore, increased PGRN triggered lysosome biogenesis by regulating transcription factor EB and accelerated mitophagy flux in VSMC, while treatment with spermidine, an autophagy inducer, improved mitochondrial phenotype and enhanced vascular contractility. Finally, angiotensin II failed to induce vascular contractility in PGRN-/- suggesting a key role of PGRN to maintain the vascular tone.

**Conclusion:** Our findings suggest that PGRN preserves the vascular contractility via regulating mitophagy flux, mitochondrial complex I activity, and redox signaling. Therefore, loss of PGRN function appears as a pivotal risk factor in CVD development.

## Introduction

Cardiovascular diseases (CVDs) encompass a broad array of conditions that impact the heart and blood vessels. In the year 2020, CVDs was responsible for an estimated 19.1 million deaths across the globe^1^. They can manifest as congenital anomalies or be acquired, and some may also have a hereditary component^2^. Vascular smooth muscle cells (VSMCs) are the predominant cell type in the arterial wall and normally adopt a quiescent, contractile phenotype to regulate vascular tone, which is a key process in CVDs. In CVDs, VSMCs lose their inherent contractile state to a synthetic phenotype, thereby resulting in functional alterations within the arterial walls and contributing to the progression of CVD^3^. Thus, therapies aimed at preventing loss of contraction are a potential strategy for treating patients with CVDs.

While the pathogenesis of CVD is multifaceted, recent findings suggest that mitochondrial dysfunction plays a central role in both the development and progression of CVD and its associated complications^4^. Mitochondrial dysfunction has been identified as a significant factor in the phenotypic transition of VSMCs^5^ and the development of arterial dysfunction^6,7^. In CVDs, mitochondrial dysfunction leads to the generation of reactive oxygen species (mtROS), decreased ATP production, thereby resulting in loss of contraction of VSMCs. However, the precise mechanisms underlying mitochondrial functions in VSMCs remain largely unknown. Hence, there is a pressing need to explore innovative therapeutic and preventive strategies focused on revitalizing VSMC mitochondrial function. This pursuit involves leveraging key components responsible for maintaining mitochondrial homeostasis, which encompasses a complex network of cellular processes including mitochondrial fission, fusion, biogenesis, and mitophagy. Disruptions in these intricate mechanisms can lead to mitochondrial dysfunction and subsequent organ damage. Notably, in the context of CVDs, deviations in mitochondrial structure and function are linked to decreased levels of mitophagy, highlighting the potential importance of mitophagy in sustaining mitochondrial homeostasis and preserving VSMC function^8,9^.

Progranulin (PGRN), a secreted glycoprotein, demonstrates widespread expression across diverse tissues and cell types, with its multifaceted roles encompassing embryogenesis, inflammation, wound healing, neurodegenerative processes, and lysosomal function^10^. Within the realm of neurodegenerative disorders, specifically frontotemporal lobar degeneration (FTLD), mutations in the granulin (GRN) gene result in PGRN protein haploinsufficiency, precipitating neurodegeneration^11^. In the cardiorenal domain, PGRN has been identified as a guardian of cardiac and renal integrity during ischemia-reperfusion injury^12,13^. Furthermore, PGRN exhibits anti-inflammatory effects within the vascular endothelium, bestowing athero-protective benefits that include elevating nitric oxide (NO) levels^14^ and protecting the endothelium integrity and blood pressure (BP) control as demonstrated recently by our group^15^. In VSMC, PGRN acts as a mitigator of calcification^16^ and regulator of cell migration via modulating IL-8 secretion^17^. Although the vascular protective effects of PGRN have been described before in other scenarios, it is still unclear if it regulates vascular contractility.

In this study, we unveil a novel facet of PGRN’s function, wherein deficiency in PGRN levels induce loss of VSMC contractility. Crucially, our novel findings uncover an innovative role for PGRN in preserving mitochondrial equilibrium through mtROS signaling, mitochondrial complex I biogenesis and function, and mitophagy flux in VSMCs. This discovery implies that PGRN may present a highly promising therapeutic avenue for tackling cardiovascular diseases by modulating mitochondrial function and cellular bioenergetics and further suggest that individuals with PGRN deficiency display cardiovascular risk along with neurodegenerative diseases.

## Material and methods

### Mice

Twelve- to sixteen-week-old male and female C57BL6/J wild type (PGRN+/+), global PGRN mutant B6(Cg)-Grn^tm1.1Aidi^/J (PGRN-/-), and male mt-Keima mice were used. All mice were fed with standard mouse chow and tap water was provided ad libitum. Mice were housed in an American Association of Laboratory Animal Care–approved animal care facility in the Rangos Research Building at the Children’s Hospital of Pittsburgh of the University of Pittsburgh. The Institutional Animal Care and Use Committee approved all protocols (IACUC protocols # 19065333 and 22061179). All experiments were performed in accordance with Guide Laboratory Animals for The Care and Use of Laboratory Animals.

### Lentivirus encoding PGRN delivery in fresh aortae

Lentiviral particles (LV) were packaged and transfected by Vectorbuider as pLV[Exp]-Puro-CMV>mGrn[NM_008175.5] with mPGK promotor. The thoracic aortae from both PGRN+/+ and PGRN-/- mice were aseptically isolated and subsequently rinsed with PBS. Following this, they were placed in individual wells of a 96-well plate, each containing 100 µL of Dulbecco’s Modified Eagle Medium (DMEM). The aortae were then categorized into two groups: control and treatment with LV. In the treatment wells, 1 µL of a 10^6^ LV vector was added. The entire culture plate was placed in a 37°C incubator with a 5% CO2 atmosphere for a duration of 24 hours.

After this 24-hour incubation period, the aortae were again washed with PBS to remove any residual substances. They were subsequently employed for various analyses, including western blot analysis, myography, and oxygen consumption rate (OCR) experiments.

### Vascular function

Thoracic aortic rings were utilized in our experiments and mounted in a wire myograph (Danysh MyoTechnology) for isometric tension recordings, employing PowerLab software (AD Instruments) as previously described in the literature^18-20^. Briefly, rings measuring 2mm in diameter were dissected from thoracic aorta and the endothelium was removed by using a needle 25G. Aortic rings were carefully placed in tissue baths containing Krebs Henseleit Solution, which was maintained at a temperature of 37 °C and continuously aerated with a mixture of 95% O2 and 5% CO2. The composition of the Krebs Henseleit Solution was as follows (in mM): 130 NaCl, 4.7 KCl, 1.17 MgSO4, 0.03 EDTA, 1.6 CaCl2, 14.9 NaHCO3, 1.18 KH2PO4, and 5.5 glucose.

### Protocol to study if deficiency in the PGRN affects vascular contractility

KCL (120mM)-induced contractility and concentration response curve (CRC) to phenylephrine (PE, alpha-1 adrenergic receptor-dependent vasoconstrictor) and U46619 [thromboxane A2 (TXA2) mimetic] were analyzed in endothelium-denuded rings from thoracic aorta from PGRN+/+ and PGRN-/- mice.

### Ex vivo protocol for regain of PGRN function in isolated aortae

Aortae were harvested from PGRN+/+ and PGRN-/- mice, the endothelium was removed, rings (2 mm) were isolated, and endothelium was removed as described above. Then, rings were incubated with LV encoding PGRN (10^6^/mL) for 24 hours in DMEM containing 100 U/ml penicillin, 100 μg/ml streptomycin, and 10 mmol/L Hepes. Reexpression of PGRN in aortae from PGRN-/- or overexpression of PGRN in aorta from PGRN+/+ were confirmed by western blot.

### High-Resolution Respirometry

To investigate the impact of PGRN on mitochondrial function, we used Oroboros Oxygraph-2K (Oroboros, Österreich, Austria) ^21^. Aortas from PGRN+/+ and PGRN-/- were evaluated using an were were freshly isolated, carefully dissected, weighed, and placed into 500 μL of isolation buffer composed of 225 mM mannitol, 75 mM sucrose, 10 mM Tris pH 7.4, and 0.2 mM EDTA. The aortas were then transferred to O2-equilibrated chambers containing 5 mL of isolation buffer. Once the baseline became stable, 10 μM cytochrome C (Sigma, St. Louis, MO) was introduced to assess mitochondrial integrity.

To evaluate mitochondrial respiration, a series of substrates and inhibitors were added sequentially. Specifically, malate (5 mM, Sigma, St. Louis, MO), ADP (2 mM, Sigma-Aldrich Co., St. Louis, MO), pyruvate (5 mM, Sigma-Aldrich Co.), glutamate (5 mM, Sigma-Aldrich Co.), and succinate (5 mM, Sigma-Aldrich Co.) were successively administered to stimulate State 3 respiration. State 3 respiration is often used in research to assess the capacity of mitochondria to produce ATP under conditions of high energy demand^22,23^. To uncouple the electron transport chain (ETC), carbonyl cyanide m-chlorophenyl hydrazone (CCCP) (1mM, Sigma-Aldrich Co.) was added. Finally, to inhibit complex I, Rotenone (1mM, Sigma, St. Louis, MO) was introduced to halt mitochondrial respiration entirely^24^.

### Vascular remodeling

Mice were euthanized to obtain aortae, followed by perfusion with chilled phosphate-buffered saline (PBS). Harvested aortae were then immersed in a 4% paraformaldehyde (PFA) solution for histological examination. After 12 hours in PFA, the tissues were transferred to 70% ethanol, where they remained until sample preparation for histology. The aortae were embedded in paraffin, after which the samples were sectioned and subjected to hematoxylin and eosin (H&E) and Masson’s trichrome staining to evaluate vascular remodeling and structural characteristics.

### Pharmacological interventions in mice

### Restoration of circulating PGRN

To restore circulating PGRN levels, PGRN+/+ and PGRN-/- mice were subjected to a 7-day treatment regimen using rPGRN delivered through ALZET osmotic minipumps (Alzet Model 1001; Alzet Corp Durect, Cupertino, CA) at a rate of 20ug/day, following established protocol by our group(*15*). After this 7-day treatment period, we conducted the analysis of mitochondrial respiration employing Oroboros O2k respirometry.

### Autophagy inducer

Spermidine, recognized as a natural polyamine with autophagy-stimulating properties^25^ was employed in this study. Both PGRN+/+ and PGRN-/- mice were randomly allocated into two cohorts: the spermidine treatment group and the control group. The treatment group received a 14-day regimen of spermidine administered via their drinking water at a concentration of 3mM. After this 14-day treatment period, evaluations were conducted, including the assessment of vasocontraction responses in thoracic aortas and the analysis of mitochondrial respiration employing Oroboros O2k respirometry. In addition, hearts were harvested for further analysis of autophagy marker through western blotting.

### Angiotensin II infusion

Both PGRN+/+ and PGRN-/- mice were subjected to a 14-day infusion protocol using ALZET osmotic minipumps (Alzet Model 1002; Alzet Corp Durect, Cupertino, CA) to delivery Angiotensin-II (Ang-II) at a dose of 490 ng/min/kg ^15,26^. Subsequently, thoracic aortas were utilized for a series of experiments, including assessments of vascular reactivity, histological analyses, and mitochondrial respiration.

### Fresh primary VSMC isolation from thoracic aorta

We conducted the isolation of VSMC from the aortas of male PGRN+/+ and PGRN-/- mice. The isolation procedure followed a well-established enzymatic dissociation protocol^27-30^. Following the isolation, the aortic smooth muscle cells were cultured in DMEM from Invitrogen Life Technologies. To maintain cell health and preserve their physiological characteristics, the culture medium was supplemented with 10% fetal bovine serum (FBS) obtained from HyClone, along with 100 U/ml penicillin, 100 μg/ml streptomycin, and 10 mmol/L Hepes (pH 7.4) from Sigma-Aldrich. To ensure the viability and functionality of the arterial smooth muscle cells during experimentation, we utilized cells within passages 4 to 8 ^31^.

### Rat Aortic Smooth Muscle cells

Rat Aortic Smooth Muscle cells (RASMC) (Lonza, Walkersville, MD) and meticulously maintained in DMEM (Gibco from Thermo Fisher Scientific, Waltham, MA, U.S.A) supplemented with 10% FBS (HyClone Logan UT, U.S.A), 100 U/ml penicillin, and 100 μg/ml streptomycin (Gibco from Thermo Fisher Scientific, Waltham, MA, U.S.A) as described before^32^.

### VSMC overexpressing PGRN via lentiviral transduction in rat aortic smooth muscle cells (aSMC)

Lentiviral particles (LV) were packaged and transfected by Vectorbuider as pLV[Exp]-Puro-CMV>mGrn[NM_008175.5] with mPGK promotor. One day prior to infection, 1×10^5^−2×10^5^ selected aSMC per well were seeded in a six-well plate, ensuring they reach an approximate confluency of 80% before initiating infection. On Day 1, the pre-existing media was aspirated, and each well was supplemented with 400 ul of fresh media, along with 5 ul of Polybrene at a concentration of 8 μg/mL as a transfection enhancer. Furthermore, 1 ul of LV containing 1x10^6^ viral particles is added to each of the wells. Control wells, one containing only cells and the other with media alone, were also maintained. The cells were subsequently incubated for a period of 48 to 72 hours without changing the media. On Day 3, the media was exchanged with 2 ml of DMEM supplemented with Puromycin at a concentration of 10 ug/ml. The introduction of Puromycin is delayed by at least 24 hours post-infection to ensure adequate expression of the Puromycin resistance gene. Day 4 entails the removal of media, which was preserved at -20°C for future use, followed by its replacement with fresh DMEM containing Puromycin at a concentration of 5 ug/ml. Finally, on Day 5, the media is once again replaced with DMEM containing Puromycin at 5 ug/ml. Throughout this procedure, special attention was directed towards monitoring the positive control well, as it had the potential to exhibit cell death in contrast to the LV-infected cells. The cells were then washed with phosphate-buffered saline (PBS) and incubated in regular medium for 48 h, after which PGRN expression was confirmed by western blot in cell lysate and ELISA (R&D biosystem) in the supernatant of SMC (control) and SMC^PGRN^ (high expression of PGRN). All cell culture processes were diligently conducted under conditions of 5% CO2 at 37°C, with the experiments thoughtfully executed in triplicate to ensure robustness and reliability of the results.

### Pharmacological treatments in cells

In our experimental setup, we employed specific treatments to investigate the autophagy pathway, mitochondrial ROS, and detrimental pathways. Bafilomycin (50 μM), was used as an inhibitor of autophagosome-lysosome fusion to confirm the ability of PGRN to induce autophagy. Recombinant PGRN (rPGRN) (600ng/mL) was used to examine if circulating PGRN regulates mitochondria respiration. Cells were also treated with Platelet-Derived Growth Factor-BB (PDGF) (100 ng/ml) to analyze where PGRN deficient cells are more sensitive to a deleterious environment. Antimycin A (20 μM) was used to induce mtROS formation.

### Collagen contraction assay

To assess the contractility of VSMC, a collagen gel contraction assay was conducted^33^. In the gel preparation process, 50 μL of 100 mM/L NaOH was added to 400 μL of rat tail collagen (PurColl, Advanced Biomatrix, #5005) kept on ice. This was followed by the addition of 50 μL of 10×PBS. Subsequently, 280 μL of VSMCs suspension (1.2×105 cells/ML) was introduced to form the gel-cell mixture, which was then transferred to the wells of a 12-well plate and allowed to further incubate for 30 minutes at 37 ℃^34^. A negative control lacking cells was also established. After gelation, 500 μL of culture medium was added to each well. The VSMC/collagen gel was left to float freely by gently scraping with a cell scraper. Contraction extent was evaluated by measuring gel area and diameter immediately (baseline) and every 24 hours using a dissection microscope. The plates were then placed back in an incubator at 5% CO2 and 95% air. All experiments were carried out in triplicate over a 72-hour period. For data analysis, the ImageJ imaging analysis software (National Institutes of Health, Bethesda, MD) was employed to measure the gel area. The final data was presented as mean ± SEM with triplicate wells in each group, and the data was reported in terms of area in mm².

### Phalloidin fluorescence

Phalloidin fluorescence was employed for visualizing the cellular cytoskeleton^32^. Cells were cultured on coverslips overnight. Subsequently, they were fixed using a 3.7% formaldehyde solution in PBS for 15 minutes at room temperature. Permeabilization was accomplished by treating the cells with 0.1% Triton™ X-100 in PBS for 15 minutes. A blocking step followed, involving a solution containing 1% BSA for 45 minutes at room temperature. The cells were then exposed to a phalloidin staining solution for 60 minutes at room temperature. Throughout the process, residual solutions were removed through PBS washes. DAPI staining was employed for visualizing cell nuclei. Microscopic imaging was conducted using a fluorescence microscope (Revolve, Echo, San Diego, California, USA).

### Seahorse Extracellular Flux Assay

The sensor cartridge and utility plate (Agilent, Santa Clara, CA) were hydrated and prepared according to company protocol. Primary VSMCs (2x10^4^-3x10^4^) were seeded onto to an Xfe96 poly-D-Lysine (PDL) Cell Culture Plate (Agilent) in 80uL of 5% FBS, 1% pen/strep complete DMEM (Sigma, St. Louis, MO) and incubated for 1 hr at room temperature for cell adhesion. Complete DMEM was removed and replaced with 180uL of Xfe DMEM Medium, pH 7.4 (Agilent) supplemented with 25mM D-(+)-Glucose (Sigma), 1mM Sodium Pyruvate (Sigma), and 2mM L-Glutamine (Corning, Corning, NY) and incubated in a non-CO2 37oC incubator for an additional 1 hr. Seahorse XF Cell Mito Stress Test Kit (Agilent, Santa Clara, CA) Oligomycin (Oli), carbonyl cyanide-p-trifluoromethoxy phenylhydrazone (FCCP), and Rotenone(ROT) reagents were reconstituted in 420 uL and 720 uL of XF Assay Media (Agilent), respectively. Each reagent was loaded into its respective port in sensor cartridge for a final well concentration of 1.0 uM Oli (ATP synthase inhibitor), 1.0 uM FCCP (uncoupling agent), and 0.5 uM ROT (complex I inhibitor)^35^. The sensor cartridge was loaded into XF96 Seahorse Extracellular Flux Analyzer (Agilent) for sensor calibration followed by addition of cell culture plate. The data were analyzed using Xfe software 2.6.1 (Agilent Technologies, Inc.) and normalized to the amount of protein loaded per well.

### Real-Time Polymerase Chain Reaction (RT-PCR)

mRNA extraction from aortae and VSMC was accomplished using the Rneasy Mini Kit (Qiagen, Germantown, MD, USA). To synthesize complementary DNA (cDNA), reverse transcription polymerase chain reaction (RT-PCR) was performed utilizing SuperScript III (Thermo Fisher, Waltham, MA, USA). The reverse transcription process occurred at 58 °C for 50 minutes, succeeded by enzyme inactivation at 85 °C for 5 minutes. For real-time quantitative RT-PCR, the PowerTrack™ SYBR Green Master Mix (Thermo Fisher, Waltham, MA, USA) was employed. The gene sequences utilized are provided in the Supplementary Table 1. Experiments were conducted on a QuantStudio™ 5 Real-Time PCR System using a 384-well format (Thermo Fisher, Waltham, MA, USA). Data analysis was executed using the 2ΔΔCt method, with results represented as fold changes signifying either upregulation or downregulation.

### Immunoblotting (Western Blot)

Aortic protein extraction was carried out using a radioimmunoprecipitation assay (RIPA) buffer (Thermo Fisher Scientific Inc.), supplemented with protease inhibitor cocktail (Roche), which consisted of 30 mM HEPES (pH 7.4), 150 mM NaCl, 1% Nonidet P-40, 0.5% sodium deoxycholate, 0.1% sodium dodecyl sulfate, 5 mM EDTA, 1 mM NaV04, 50 mM NaF, 1 mM PMSF, 10% pepstatin A, 10 μg/mL leupeptin, and 10 μg/mL aprotinin. Total protein extracts obtained from aortic homogenates were subjected to centrifugation at 15,000 rpm for 10 minutes, and the resulting pellet was discarded, 25 μg of protein was used. For aVSMC and rat aSMC samples, direct homogenization was performed using 2x Laemmli Sample Buffer supplemented with 2-Mercaptoethanol (β-mercaptoethanol). Protein samples (were separated by electrophoresis on a polyacrylamide gradient gel, and subsequently transferred to a PVDF membrane (Immobilon FL, EMD Millipore, Billerica, MA). To prevent non-specific binding, the membranes were blocked with either 5% skim milk or 1% bovine serum albumin (BSA) in tris-buffered saline solution with tween for 1 hour at 24 °C. Specific antibodies, as listed in supplementary table 2, were then applied to the membranes, and incubated overnight at 4 °C. Following the primary antibody incubation, secondary antibodies were used, and the membranes were subjected to enhanced chemiluminescence with a luminol reagent (SuperSignal™ West Femto Maximum Sensitivity Substrate, Thermo Fisher Waltham, MA, USA) was used for antibody detection.

### Mitochondrial related experiments Complexes expression-OXPHOS

We employed the OXPHOS cocktail antibody mixture (Abcam), specifically designed to target complexes subunits within the Oxidative Phosphorylation (OXPHOS). To ensure the preservation of these hydrophobic protein’s integrity, a meticulous methodology was adhered to. Prior to initiation, cells were collected and lysed for protein extraction, with utmost care taken to maintain the integrity of OXPHOS complexes. Samples were strictly kept unheated before their application onto the gel.

### Mitochondrial complex I activity

The assessment of mitochondrial respiratory chain complex I activity was conducted by using the specialized Complex I Enzyme Activity Assay Kit (ab109721). Initially, cells were lysed, and extracts from these cells were carefully loaded onto a microplate. Subsequently, the plate wells were meticulously washed three times with a designated buffer. Following this preparation, 200 µL of assay solution was added to each well, and the optical density (OD450 nm) was measured in kinetic mode at room temperature for up to 30 minutes (SpectraMax i3x Multi-Mode Microplate Reader). Absorbance was analyzed at OD 450 nm, which was used as an indicator of Complex I activity.

### NAD+/NADH assay

The NAD+/NADH ratio was assessed using the NAD+/NADH Assay Kit (ab65348). Initially, cells were digested in 120 μL of NADH/NAD extraction buffer. After centrifugation in a 10 kD Spin Column (ab93349) at 14,000 x g for 20 minutes at 4°C, half of the sample was transferred to a new tube and incubated at 60°C for 30 minutes to decompose NAD+, while the remaining half was designated as NADt (comprising both NADH and NAD+). Subsequently, 20 μL of NADt and 20 μL of the decomposed NAD+ sample were mixed with 30 μL of extraction buffer and then incubated with 100 μL of reaction mix at room temperature for 5 minutes, facilitating the conversion of NAD+ to NADH. Following this, 10 μL of NADH Developer was added to each well, and the mixture was allowed to react at room temperature for 20 minutes. The resulting sample outputs were measured at OD 450 nm on a microplate reader (SpectraMax i3x Multi-Mode Microplate Reader) in kinetic mode.

### Measurement of ATP concentration

ATP concentration of the cells was determined by ATP Assay Kit (ab83355) according to the manufacturer’s protocol. Cells were seeded at 1× 10^6^ cells/mL. The cells were collected and lysed in 150 µL of ATP assay buffer. Then, the samples were centrifuged for 5 min at 4 °C at 13,000× *g* to remove insoluble material and loaded to a 96-well plate (ThermoFisher Scientific) at a volume of 50 µL in 3 technical replies. The plate was incubated at room temperature for 30 min and protected from light. The absorbance was measured at wavelength λ = 570 nm by the spectrophotometer (SpectraMax i3x Multi-Mode Microplate Reader). The experiment was conducted using four biological replications and the collected data were calculated according to the producer’s instructions. The data were presented as an ATP concentration (nmol) per 1 mg of total protein in each well.

### Mitochondrial membrane potential

For the analysis of mitochondrial membrane potential (MMP), cells were subjected to JC-1 working solution (10 μg/ml) and subsequently incubated in a 5% CO2 incubator at 37 °C for a duration of 30 minutes. Following this incubation period, the JC-1-treated cells underwent three washes with PBS, with each wash lasting 2 minutes. Images were then acquired using a fluorescence microscope (Revolve, Echo, San Diego, California, USA). Carbonyl Cyanide Chlorophenylhydrazone (CCCP, 50uM for 24h)

### Mitochondrial ROS production

### MitoSOX-based flow cytometric assay

To evaluate mitochondrial ROS levels, we employed a MitoSOX-based flow cytometric assay on both control cells and cells subjected to antimycin A treatment (20uM for 30 minutes). Initially, primary VSMC from PGRN+/+ and PGRN-/- or SMC and SMC^PGRN^ were seeded in six-well plates and subsequently exposed to MitoSOX (5 μM) staining in complete medium for a half-hour at 37 °C. After staining, a gentle wash followed by a 5-minute trypsinization at 37 °C was performed. Subsequently, the cells were resuspended in FACS buffer (comprising PBS, 1% BSA, and 1 mM EDTA) and subjected to immediate analysis via flow cytometry. The flow cytometer recorded both mean fluorescence intensity and the percentage of stained cells. The data were presented as histograms representing the cell count emitting specific fluorescence, effectively illustrating the size of the corresponding cell population. Graphs were generated to depict the mean fluorescence intensity, offering a quantifiable representation of the observed results. Additionally, the effects of antimycin A were presented by delta (difference between before and after drug incubation) in primary and cells overexpressing PGRN.

### Lucigenin chemiluminescence assay

The chemiluminescent probe lucigenin (bis-N-methylacridinium nitrate) was used to assess ROS levels within the cells. A cell suspension, comprising up to 1 × 10^6 cells in 1 ml of Complete Phosphate-Buffered Saline (CPBS), was supplemented with 175 µL of lucigenin (0.005 mmol/L) and assay buffer. This mixture was then meticulously transferred to a White 96-well microplate. Subsequently, the microplate was promptly inserted into a luminometer (FlexSation 3 microplate reader, Molecular Devices, San Jose, USA) to record the chemiluminescent response. These measurements were conducted at a constant temperature of 37°C and spanned a duration of 60 minutes. Following the initial reading, NADPH (0.1 mmol/L), serving as a substrate for the NADPH oxidase enzyme, was introduced to the suspension. A second reading of the microplate was subsequently conducted. The results were expressed as relative light units (RLU) per protein levels. To confirm the mitochondrial origin of the ROS signal, VSMC samples underwent incubation with antimycin A for 1 hour.

### Extraction and Quantification of Mitochondrial DNA Content and mitochondrial biogenesis

Relative mtDNA copy numbers were estimated by real-time quantitative PCR according to previously described methods^36,37^. Total DNA was isolated from cells adherent cultures by Genomic Mini Kit (Qiagen Dneasy blood & tissue kit) according to the producer’s instructions. DNA probes were diluted in Rnase-free H_2_O (Sigma-Aldrich), to equal concentration and stored at −20 °C until further use. The Mitochondrial DNA (mtDNA) content was assessed by absolute quantification of DNA copy numbers using real time PCR. mtDNA content analysis was performed with TaqMan Master Mix (Applied Biosystems) on Applied Biosystems^®^ 7500 Real-Time PCR System (Life Technologies). mtDNA was detected using primers MT16520F and MT35R, in the presence of probe MT16557TM. Nuclear DNA content was estimated by amplification of a fragment from the single-copy gene Kir4.1, with primers KIR835F and KIR903R used together with probe KIR857TM. The results were calculated as mtDNA/nDNA content based on the ΔΔCt method according to the previously described method^38^. Peroxisome proliferator-activated receptor-γ coactivator-1α (PGC-1α), a transcription factor controlling many aspects of oxidative metabolism including biogenesis^39^, was analyzed via western blot.

### Mitochondrial morphology and dynamics

Via western blot, we measured Mitofusin 1 (Mfn1) and Optic Atrophy 1 (OPA1) as markers of mitochondria fusion, and Dynamin-Related Protein 1 (Drp1) as a marker of mitochondria fission.

### Mitophagy study

To investigate mitophagy within cells, we conducted a comprehensive examination of protein expression using western blot analysis. Specifically, we assessed the levels of key markers involved in mitophagy, including LC3A, p62, PINK1, and PARKIN. To investigate mitophagy in fresh aorta, we utilized a unique transgenic mouse known as the mt-Keima mouse, graciously provided by Dr. Finkel from the University of Pittsburgh. This model is characterized by the expression of mt-Keima, a pH-dependent fluorescent protein, within the mitochondria. Mt-Keima, a coral-derived protein, is a dual-excitation ratiometric fluorescent protein known for its pH sensitivity and resilience against lysosomal proteases. In the mitochondria’s physiological pH environment (pH 8.0), it predominantly emits shorter-wavelength excitation, rendering a green appearance. However, during mitophagy within the acidic lysosome (pH 4.5), mt-Keima progressively transitions towards longer-wavelength excitation, resulting in a red fluorescence emission^40^. Following the dissection of aortae from mt-Keima mice, they were divided into control and mt-Keima-treated groups after incubation with LV for 24 hours. Subsequently, the aortae were longitudinally opened and had their endothelium removed before being mounted on glass slides and covered with coverslips. Images were acquired using a fluorescence microscope (Revolve, Echo, San Diego, California, USA).

### Lysosome biogenesis

In whole cell lysate, we studied lysosome biogenesis by measuring lysosomal-associated membrane protein (LAMP1) expression and the phosphorylation status of transcription factor EB (TFEB). In cytoplasm and nuclear fractions, we evaluated the TFEB content via western blot analysis. Fractioning was performed as described below.

### Cytoplasm and nuclear fractions preparation

Nuclear protein extracts were prepared using a nuclear extraction kit (Thermo Fisher, #78833) following the manufacturer’s guidelines. The quantification of total TFEB in cytoplasm and nuclear fractions was conducted through western blotting. For cytoplasmic fractions, GAPDH expression served as an internal control for the cytoplasm fraction, while histone3 was used as an internal control for the nuclear fractions.

### Statistic

The calculation of the maximal effect (Emax) from KCl, PE, and TXA2 were conducted through the analysis of CRC. pD2 resulting from CRCs was also calculated. CRCs were fitted using a non-linear interactive fitting program (Graph Pad Prism 9.0; GraphPad Software Inc., San Diego, CA, USA). The term “Emax” means the highest attainable effect produced by KCl, PE, and TXA2. While “pD2” indicates the negative logarithm of the molar concentration of phenylephrine and thromboxane needed to activate 50% of the maximum effect (EC50).

For comparisons of multiple groups, one-way or two-way analysis of variance (ANOVA), followed by the Tukey post-test was used. Differences between the two groups were determined using Student’s t-test. The vascular function data are expressed as a maximal response. The concentration–response curves were fitted by nonlinear regression analysis. Maximal response was determined and used to determine if there was difference between the groups. Analyses were performed using Prism 10.0 (GraphPad Software, La Jolla, CA). A difference was considered statistically significant when *P*≤0.05.

## Results

### Deficiency in PGRN affects vascular contractility

To analyze whether PGRN regulates vascular contraction, we examined the effects of KCl (120mM), phenylephrine, or thromboxane A2 analogue (U46619) in endothelium-denuded aortic rings from PGRN+/+ and PGRN-/- mice via wire myograph. Deficiency in PGRN decreased the vascular contractility to all three agents in male (Fig. 1A) and female mice (Fig. 1B) and diminished the protein expression for αSMA, but not mRNA level (Supplementary Fig. 1A and B). In addition, lack of PGRN did not affect the vascular stiffness at least in mice with 12-14 weeks of age (Supplementary Fig. 1C). Finally, deficiency in PGRN triggered vascular inflammation in male and female mice characterized by elevated ICAM, VCAM, IL1β, TNFα, and IL6 (Supplementary Fig. 1D and E).

**Figure 1.**
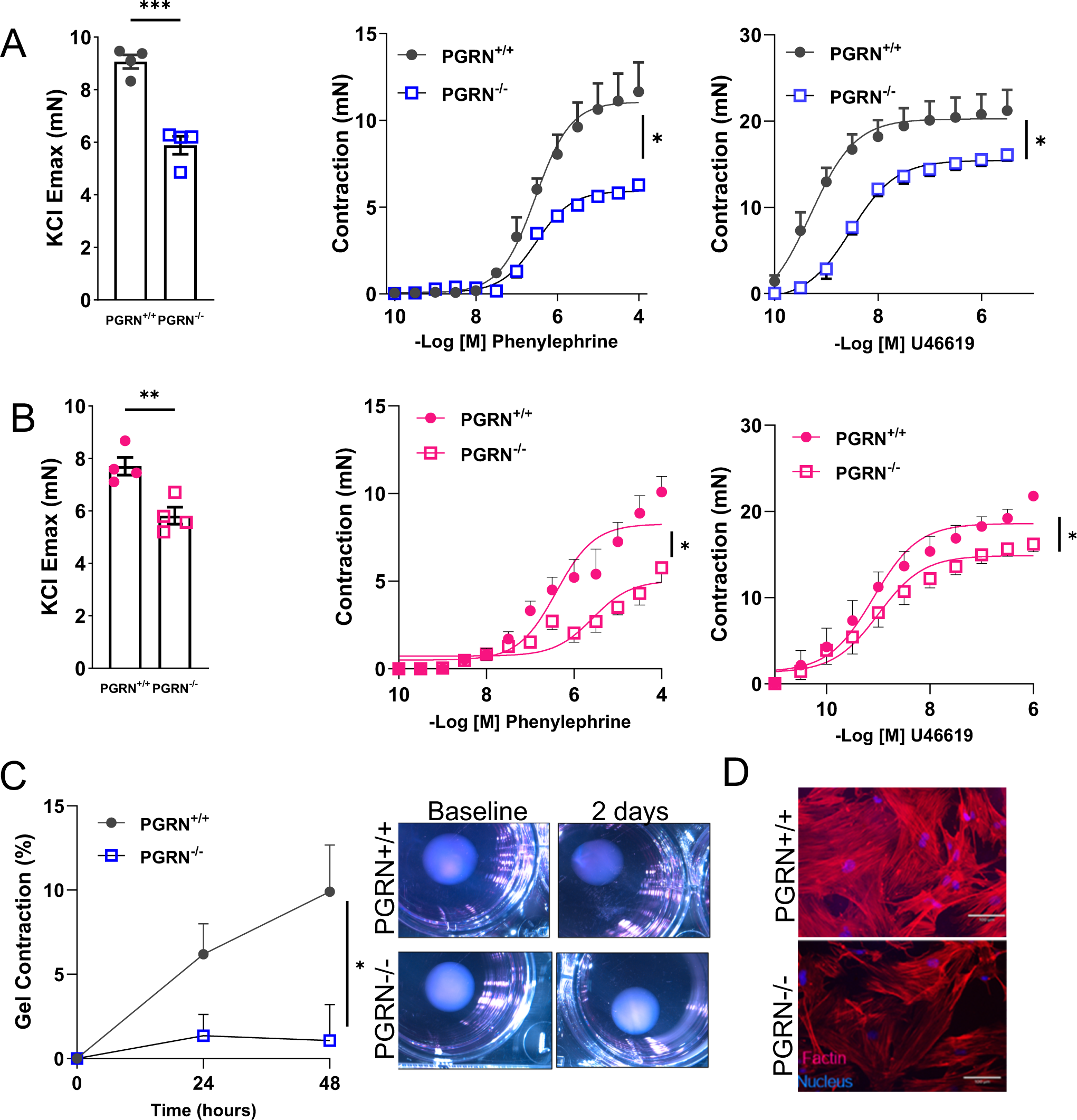
Deficiency in PGRN impairs vascular contractility *ex vivo.* *Ex vivo* wire myography showing KCl-induced vascular contractility (120mM) and concentration-effect curves to phenylephrine and thromboxane A2 analogue (U46619) in endothelium-denuded aortic rings from male (A) and female (B) PGRN-/- and PGRN+/+ mice. Collagen contraction assay in PGRN+/+ and PGRN-/- aVSMC (C). Gel area variation on day 2 post-gelation normalized to baseline area. Scale bar, 1 mm. Measurement of diameter over time. Representative image of actin filaments in PGRN+/+ and PGRN-/- aVSMC. In red, actin filaments stained with Rhodamine-Phalloidin; in blue, nuclei stained with DAPI (D). Scale bar, 100 μm. Data are presented as mean ± SEM (n = 3-6). *P < 0.05 vs. PGRN+/+ samples.

We further examined the role of PGRN regulating vascular contraction in primary VSMC isolated from PGRN+/+ and PGRN-/-. By performing the collagen gel disc assay, we observed that PGRN-/- aVSMC contracted less after 48h compared to PGRN+/+ VSMC (Fig. 1C). Furthermore, we found attenuated F-actin (Fig. 1D) and αSMA levels, with no difference in αSMA mRNA expression (Supplementary Fig. 1F and G).

### PGRN is a major regulator of mitochondria quality

In conditions like diabetic nephropathy^41^ or in neuroblastoma cells^42^, there is evidence that PGRN plays a role in the regulation of mitochondrial quality. Consequently, we sought to investigate whether PGRN also has an impact on mitochondrial profile within the vasculature. In VSMC, lack of PGRN attenuated Oxygen Consumption Rate (OCR) (Seahorse) followed by suppression of ATP levels, mitochondrial complex I activity, and NAD^+^/NADH ratio (Fig. 2A and B), but it did not affect the mitochondria number (mtDNA/nDNA) (Fig. 2C). Interestingly lack of PGRN decreased PGC1α expression (a marker of mitochondria biogenesis) (Supplementary Fig. 2A), thus suppression of PGC1α might not be affecting mitochondrial biogenesis, but other signaling instead including inflammation and antioxidant machinery^43,44^. In summary, PGRN regulates mitochondrial respiration likely adjusting complex I and without interfering with mitochondria amount, even with a suppressed PGC1α expression. Finally, mitochondria from PGRN-/- VSMC demonstrated impaired membrane potential, which was measured by JC-1 stain (Supplementary Fig. 2B). Carbonyl Cyanide Chlorophenylhydrazone (CCCP) was used as a positive control in PGRN+/+ VSMC.

**Figure 2.**
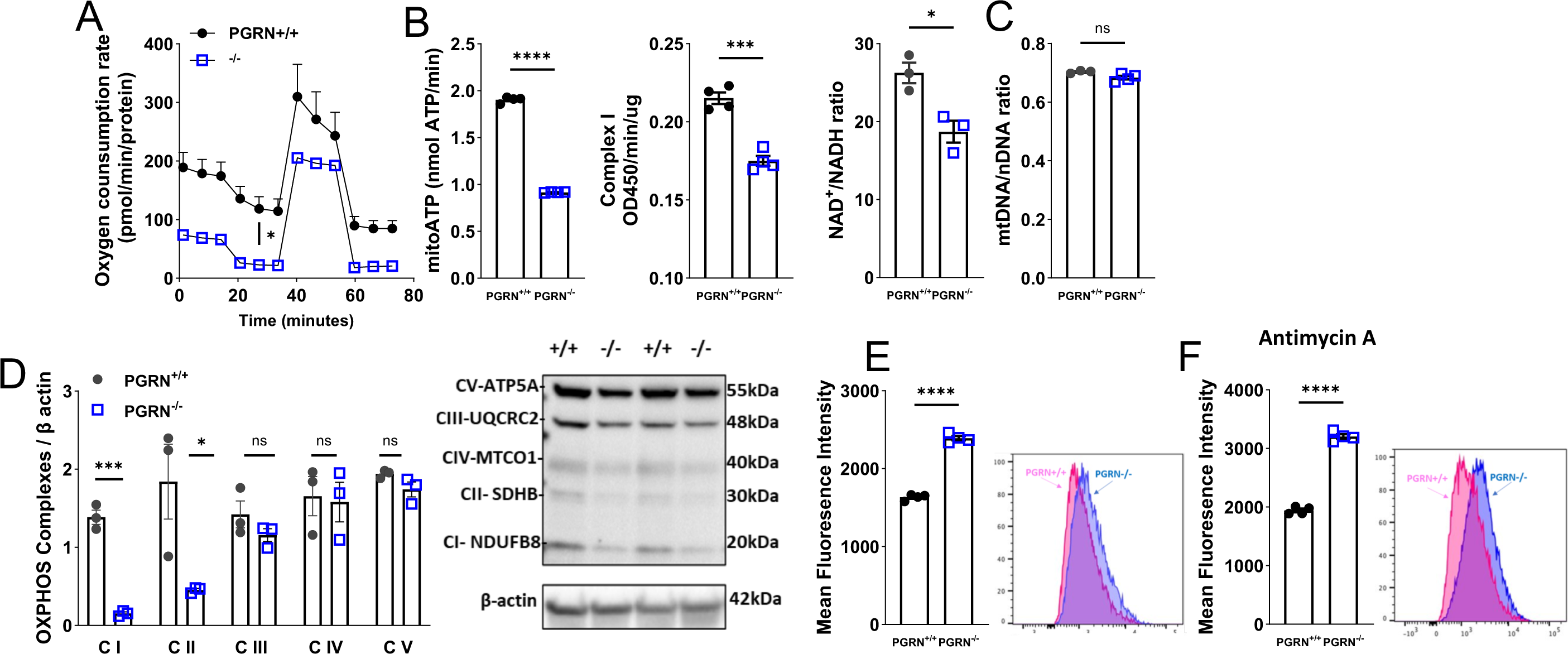
Loss of PGRN affects mitochondrial quality in aVSMC. Mitochondrial oxygen consumption rates (OCR) in PGRN+/+ and PGRN-/- aVSMC measured using the Seahorse XF Extracellular Flux 96 analyzer (A). ATP content, mitochondrial complex I activity, and NAD+/NADH ratio in PGRN+/+ and PGRN-/- aVSMC (B). Quantitative PCR (qPCR) analysis of mitochondrial DNA (mtDNA) copy number relative to nuclear DNA (nuDNA) in PGRN+/+ and PGRN-/- aVSMC (C). Western blot for protein expression of OXPHOS in PGRN+/+ and PGRN-/- aVSMCs. β-actin was used as load control (D). Mitochondrial reactive oxygen species (mtROS) levels measured by MitoSox stain via flow cytometry in PGRN-/- and PGRN+/+ aVSMCs at basal condition (E) or under Antimycin A treatment (F). Data are presented as mean ± SEM (n = 3-6). *P < 0.05 vs. PGRN+/+ aVSMC.

Since we observed suppressed complex I activity, we analyzed the complexes expression by using the OXPHOS antibody, which revealed that deficiency in PGRN reduces complexes I and II expression (Fig. 2D). The alteration of mitochondrial dynamics is essential for maintaining mitochondrial respiration. Therefore, we studied to investigate whether PGRN plays a role in orchestrating the processes of mitochondrial fusion and fission. Lack in PGRN induced mitochondria fragmentation (fission) characterized by upregulation of DRP1 expression (Supplementary Fig. 2C) and downregulated MFN1 expression (marker of fusion) (Supplementary Fig. 2D). Finally, changes in mitochondria respiration and dynamic have been associated with exacerbated mtROS formation. By using MitoSox we found that lack of PGRN induced mtROS formation (Fig. 2E), which is exacerbated by antimycin A (a potent mitochondrial ROS inducer) (Fig. 2F). Similar results were obtained by lucigenin chemiluminescence assay (Supplementary Fig. 2E). ERK1/2, a MAPK member and ROS-sensitive protein, was overactivated in PGRN-/- VSMC, which was not further affected by PDGF-BB compared to PGRN+/+ (Supplementary Fig. 2Fand G).

Additionally, we assessed whether alterations in mitochondrial function were evident in whole arteries. Using Oroboros O2k respirometry, we examined mitochondrial respiration in fresh aortae and found that mitochondrial complex I activity is impaired in PGRN-/- with no changes in complex II (coupled or uncoupled) (Fig. 3A). Furthermore, aortae from PGRN-/- displayed reduced complex I, III, and IV protein expression (Fig. 3B), complex I related genes, and peroxisome proliferator-activated receptor γ (PPARγ, a complex I gene regulator^45^) levels (Supplementary Fig. 3 and 4). Similarly, to what was observed in primary VSMC, aortae from PGRN-/- exhibited a decrease in PGC1α and MFN1 expression, along with an increase in DRP1 (Supplementary Fig. 5A-C).

**Figure 3.**
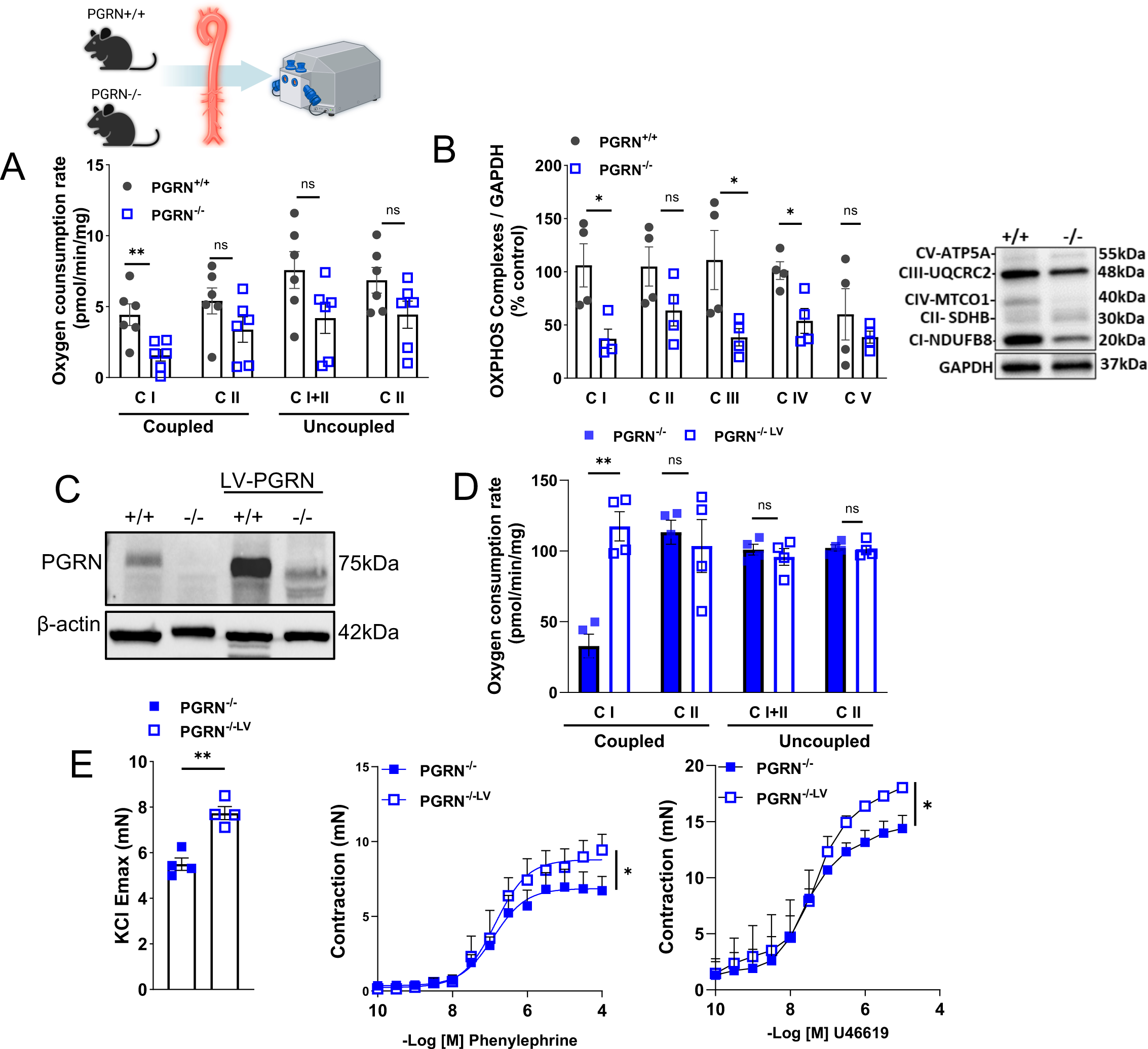
Loss of PGRN affects mitochondrial quality in fresh aortae. Re-expression of PGRN restores mitochondrial respiration and vascular contractility in PGRN deficient arteries. Mitochondrial oxygen consumption rates (OCR) in fresh aortae from PGRN+/+ and PGRN-/- measured using the Oroboros O2k respirometry (A). Western blot for protein expression of OXPHOS. GAPDH was used as load control (B). Western blot for protein expression of PGRN after lentivirus (LV) delivery for 24hours. β-actin was used as load control (C). Mitochondrial oxygen consumption rates (OCR) in fresh aortae from PGRN-/- mice after LV delivery (D). *Ex vivo* wire myography showing KCl-induced vascular contractility (120mM) and concentration-effect curves to phenylephrine and thromboxane A2 analogue (U46619) in endothelium-denuded aortic rings from male PGRN-/- mice after LV delivery. Data are presented as mean ± SEM (n = 3-4). *P < 0.05 vs. PGRN+/+ aVSMC. *P < 0.05 vs. PGRN-/-.

### Reexpression of PGRN restores mitochondrial respiration and vascular contractility in PGRN-/- deficient arteries

Circulating PGRN plays a major role in regulating cardiovascular biology. Therefore, we treated PGRN+/+ and PGRN-/- with recombinant PGRN (20ug/day/mouse) for 7 days via osmotic mini- pump^15^; we found that PGRN treatment did not affect mitochondrial respiration (Supplementary Fig. 6A). Similarly, rPGRN did not affect OCR in PGRN-/- VSMC *in vitro* (Supplementary Fig. 6B). Thus, we focused on understanding the intracellular role of PGRN by re-expressing PGRN via LV delivery. We treated the aortae from PGRN+/+ and PGRN-/- with LV encoding PGRN for 24h and analyzed the mitochondrial respiration and vascular contractility. LV increased PGRN expression in PGRN+/+ and repopulated PGRN in PGRN-/- (Figure 3C). In addition, LV increased complex I activity and vascular contractility to KCL, U46619, and phenylephrine in PGRN-/- arteries (Fig. 3D and E), but did not affect mitochondrial respiration and vascular contractility in PGRN+/+ arteries (Supplementary Fig. 7A-D)

Since we found these exciting effects of PGRN on regulating vascular contractility and mitochondrial respiration, we created VSMC overexpressing PGRN via LV delivery (aSMC^PGRN^), which was confirmed by western blot in cell lysate or ELISA in supernatant (Fig. 4A). aSMC^PGRN^ demonstrated increase in ATP levels, NAD^+^/NADPH ratio, and complex I activity (Fig. 4B). Furthermore, overexpressing PGRN reduced mtROS formation and partially protected against antimycin A-induced mtROS production (Fig. 4C). In addition, aSMC^PGRN^ demonstrated cell contraction, which could be observed in collagen gel disc assay, F-actin stain (more F-actin and smaller cell size), and higher αSMA content. (Fig. D-F). Finally, aSMC^PGRN^ demonstrated no difference for mitochondrial biogenesis and number (Supplementary Fig. 8A and B), but it increased MFN1 and diminished DRP1 expression (Supplementary Fig.8C and D).

**Figure 4.**
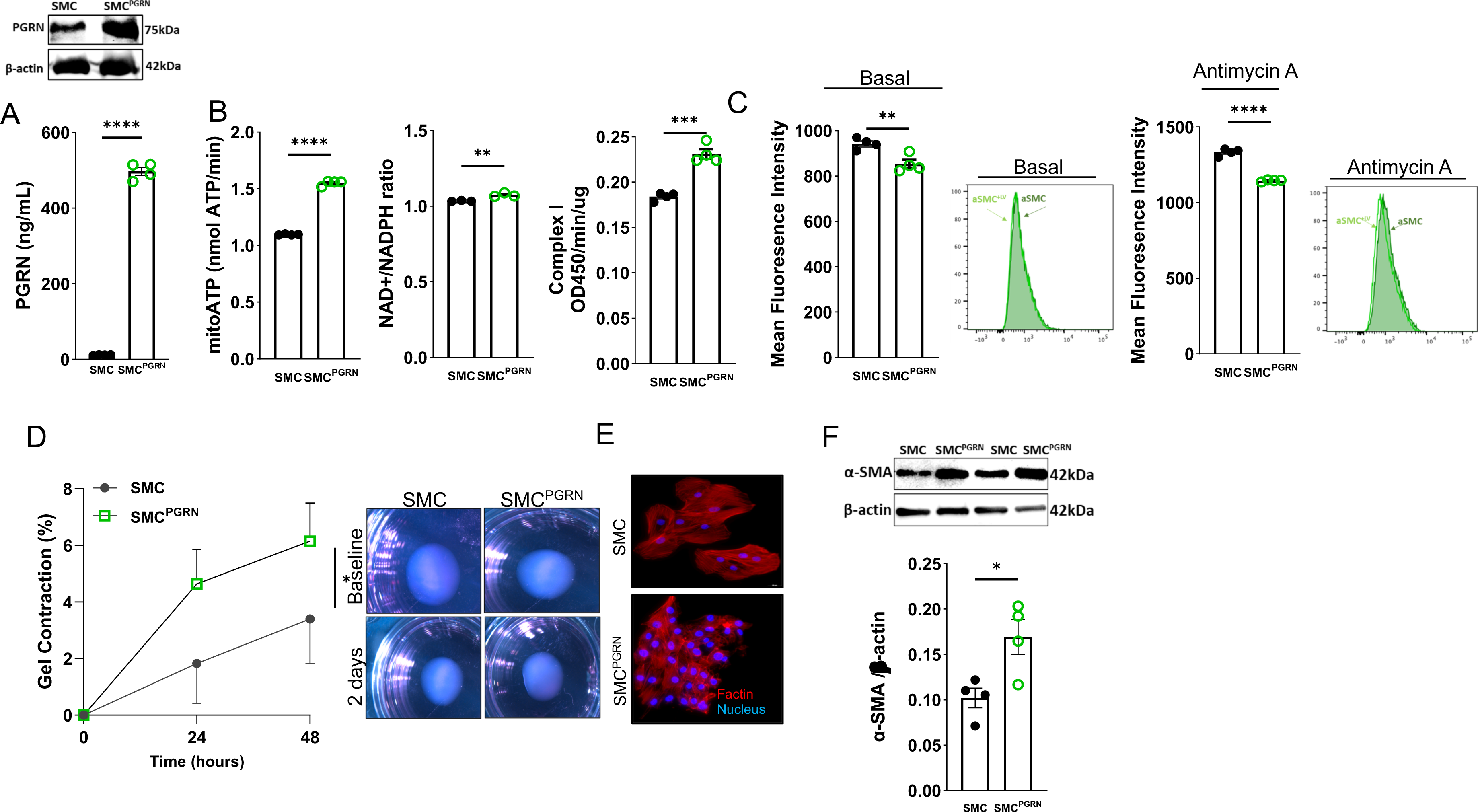
Overexpression of PGRN generates high-capacity mitochondria and induces vascular contractility in VSMC. Western blot representative for protein expression of PGRN in SMC and SMC^PGRN^ lysate or PGRN levels in the supernatant from SMC and SMC^PGRN^ β-actin was used as load control for the western blot analysis (A). ATP content, NAD+/NADH ratio, and mitochondrial complex I activity in SMC and SMC^PGRN^ (B). Mitochondrial reactive oxygen species (mtROS) levels measured by MitoSox stain via flow cytometry in SMC and SMC^PGRN^ at basal condition and under Antimycin A treatment (C). Collagen contraction assay in SMC and SMC^PGRN^. Gel area variation on day 2 post-gelation normalized to baseline area. Scale bar, 1 mm. Measurement of diameter over time (D). Representative image of actin filaments in aSMC and aSMC^PGRN^. In red, actin filaments stained with Rhodamine-Phalloidin; in blue, nuclei stained with DAPI. Scale bar, 100 μm (E). Western blot for protein expression of αSMA in aSMC and aSMC^PGRN^. β-actin was used as load control (F). Data are presented as mean ± SEM (n = 3-4). *P < 0.05 vs. aSMC. (K) Mitochondrial reactive oxygen species (mtROS) levels measured by flow cytometry in SMC and SMC^PGRN^ treated with Antimycin A. MFI: Mean fluorescence intensity.

### Deficiency in PGRN disrupts vascular mitophagy

Studies conducted on kidneys have demonstrated that PGRN regulates autophagy^41^. However, it remains unclear whether PGRN has an impact on vascular mitophagy and subsequently mitochondrial quality. Considering this, our investigation aimed to determine whether the disruption of PGRN signaling would influence mitochondrial recycling. We observed that VSMC from PGRN-/- present elevated PINK (66 and 33kDa), PARKIN, LC3I/II ratio, and p62 accumulation suggesting a dysfunctional mitophagy (Fig. 5A). Furthermore, we found an accumulation of LAMP1 (marker of lysosome), higher TFEB activity, characterized by higher TFEB content in the nuclei. (Fig. 5B). Suggesting that TFEB activation is higher in PGRN-/- VSMC, which leads to higher lysosome formation. However, despite the higher lysosome formation, they exhibit dysfunction.

**Figure 5.**
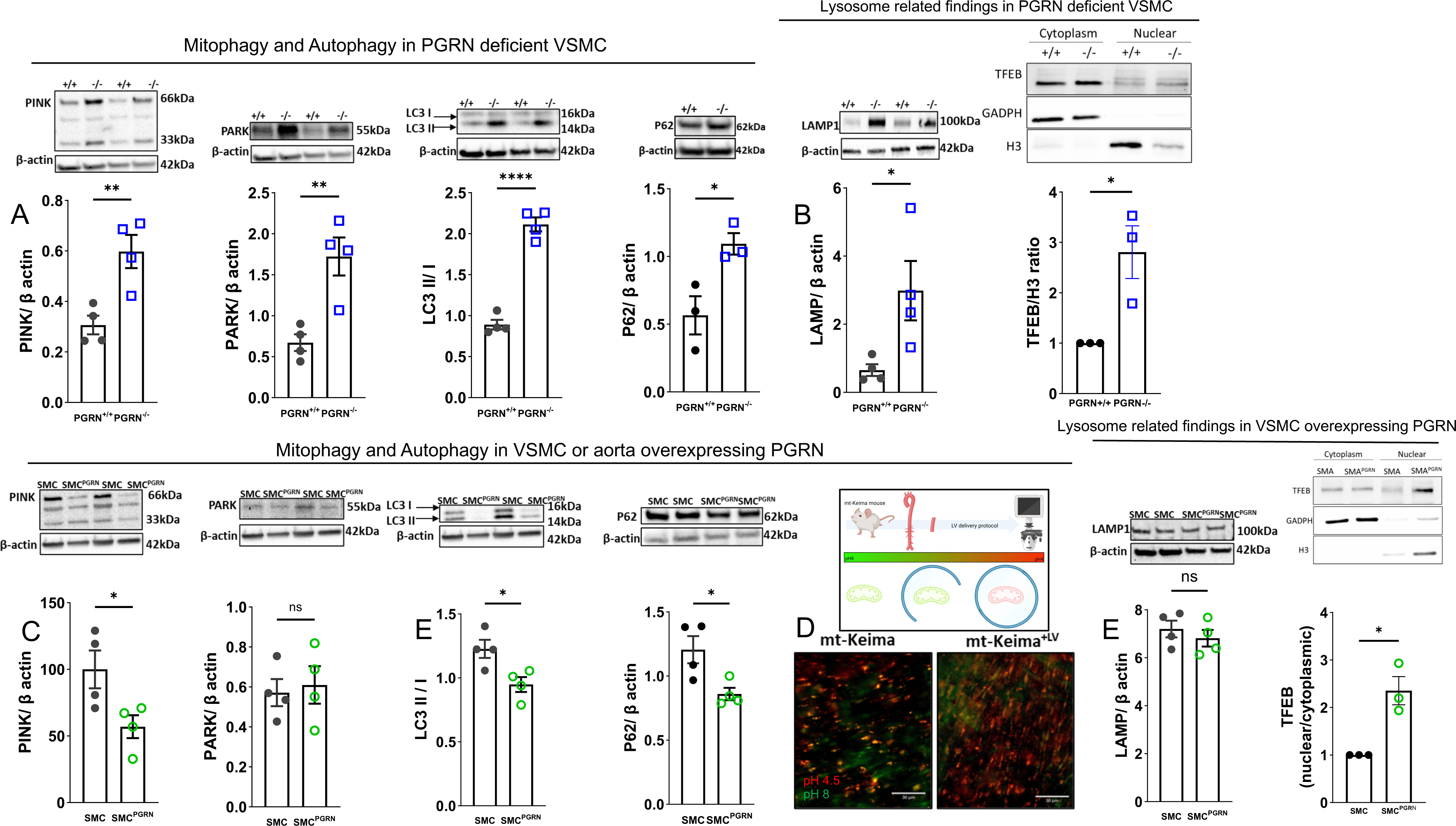
PGRN deficiency triggers dysregulated lysosome formation and disturbs mitophagy flux. Overexpression of PGRN confers an accelerated mitophagic flux in VSMC. Western blot for protein expression for mitophagy and autophagy markers - PINK (66 and 33 kDa), PARK, LC3I/II ratio, p62 (A)- lysosome levels (LAMP) and TFEB activation (B) in PGRN+/+ and PGRN-/- aVSMCs. Western blot for protein expression for mitophagy and autophagy markers - PINK (66 and 33 kDa), PARK, LC3I/II ratio, p62 (C). Track of mitophagy in fresh aortae from Mt- Keima mice treated with lentivirus (LV) encoding PGRN for 24h (D). Green represents mitochondria in pH8 (likely cytosolic), while red represents mitochondria in pH4.5 (likely lysosome). Lysosome levels and biogenesis via TFEB activation in aSMC and aSMC^PGRN^. TFEB activation was measured by quantifying TFEB content in cytosolic and nuclear fractions. GADPH was used a marker of cytoplasm, while histone3 (H3) as a marker of nuclei. β-actin was used as load control for all western blots. Data are presented as mean ± SEM (n=3-4). *P < 0.05 vs. PGRN+/+ or vs. aSMC.

In VSMC overexpressing PGRN we observed reduced PINK, LC3I/II, and p62, with no difference in PARK and LAMP (Fig. 5C and Fig.5E). Furthermore, we used arteries from mt-Keima mice to track mitophagy in freshly isolated aortae. Thus, we overexpressed PGRN via LV delivery (24h of incubation) in aortae from these mice and tracked mitochondrial flux. PGRN caused a remarkable increase of mitochondria in lysosome (red) compared to control arteries (Fig. 5D).

To confirm that high levels of PGRN induce autophagy we treated the cells with bafilomycin A1 (vacuolar H+-ATPase Inhibitor) and observed that accumulation of LC3I/II and p62 (Supplementary Fig. 9A and B) indicating that PGRN is an autophagy inducer. Finally, we found that overexpression of PGRN triggered higher presence of TFEB in the nuclei (Fig. 5E), which suggests that PGRN induces lysosome biogenesis and autophagy.

### Spermidine, an autophagy inducer, rescues mitochondrial respiration and vascular contractility in PGRN deficient mice

We used spermidine as an autophagy inducer to analyze whether restoring autophagy flux would improve mitochondrial respiration and vascular contractility (Supplementary Fig. 11A). To confirm that spermidine caused autophagy we measured LC3I/II in hearts from PGRN+/+ and PGRN-/- and observed that spermidine restored LC3A expression in PGRN-/- (Supplementary Fig. 10B). Furthermore, spermidine increased the mitochondrial respiration and vascular contractility in the aortae from PGRN-/- mice (Supplementary Fig. 10C-F). We did not see any effect of spermidine in arteries from PGRN+/+ mice (Supplementary Fig. 11A-D).

### PGRN deficient mice do not respond to Angiotensin II-induced vascular contractility

We first analyzed whether Angiotensin II changes PGRN expression in aortae from PGRN+/+, which revealed that Angiotensin II upregulated PGRN protein expression (Fig.6A). We next challenged the PGRN+/+ and PGRN-/- mice with Angiotensin II (an inducer of vascular contraction and mitochondria dysfunction) to analyze mitochondria respiration and vascular function. We found that Angiotensin II did not change the mitochondria respiration in PGRN-/- compared to PGRN+/+ (the difference between PGRN+/+ and PGRN-/- persisted) (Fig.6B) and did not affect vascular contractility in endothelium-denude aortic rings from PGRN-/- mice (Fig.6C-E), Finally, Angiotensin II similarly affected the vascular stiffness in PGRN+/+ and PGRN-/-, but PGRN-/- were more sensitive to Angiotensin II-induced vascular fibrosis (Fig. 6F). These findings imply that Angiotensin II-induced vascular contractility depends on the integrity of the PGRN signaling pathway. Additionally, it appears that PGRN plays a pivotal role as a gatekeeper in the formation of fibrosis.

**Figure 6.**
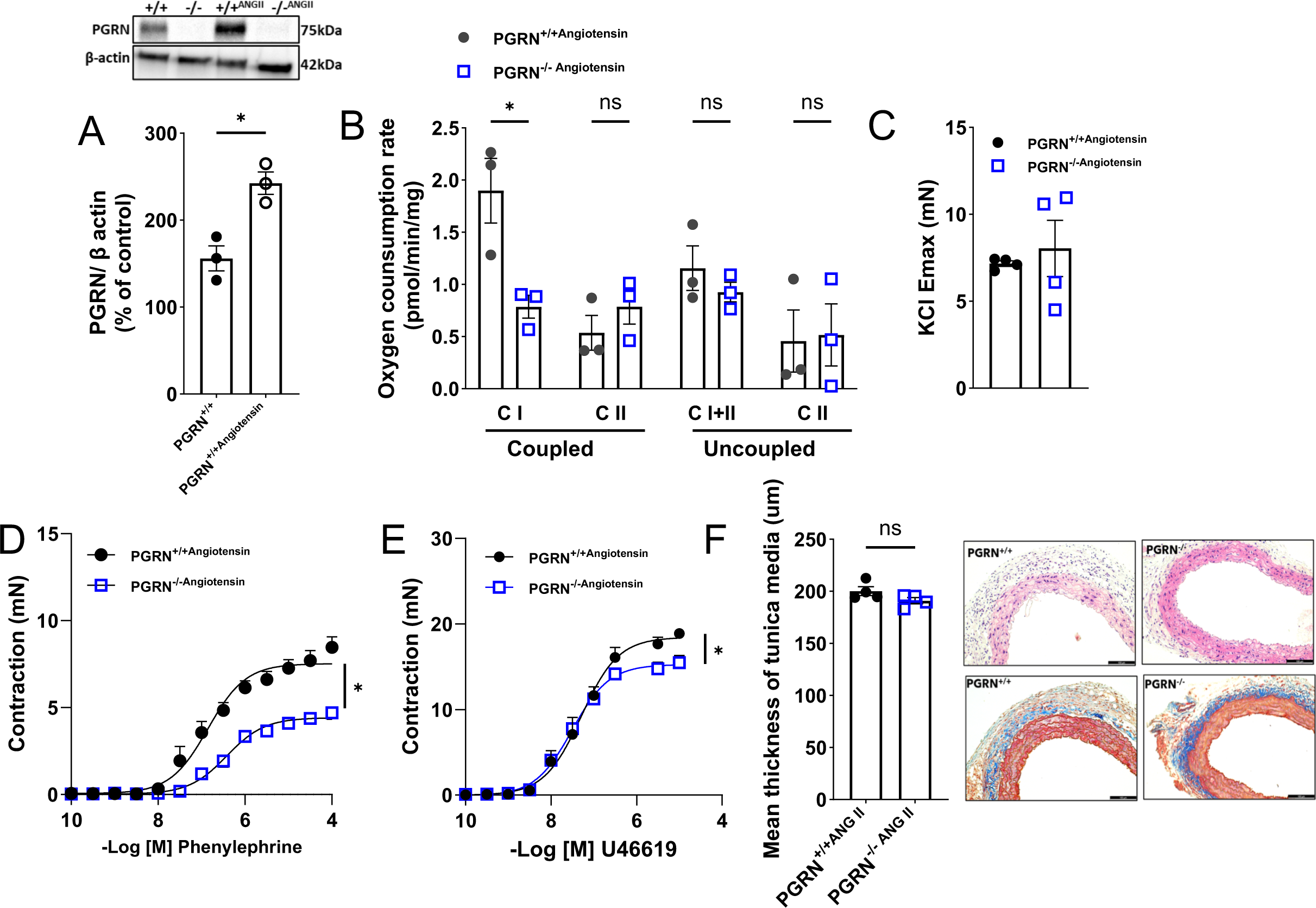
PGRN deficient mice fail to respond to Angiotensin II-induced vascular hypercontractility. Western blot for PGRN expression in aortae from PGRN+/+ and PGRN-/- mice treated with Angiotensin II β-actin was used as load control (A). Mitochondrial oxygen consumption rates (OCR) measured using the Oroboros O2k respirometry in fresh aortae of PGRN+/+ and PGRN-/- mice treated with Angiotensin II (B). *Ex vivo* wire myography showing concentration-effect curves to KCl (120mM), phenylephrine, and thromboxane A2 analogue (U46619) in endothelium-denuded aortic rings of aortae PGRN+/+ and PGRN-/- mice treated with Angiotensin II (C). Morphometric analysis of media area derived from Hematoxylin & eosin stain and Masson’s trichrome stains in aortae from PGRN+/+ and PGRN-/- mice treated with Angiotensin II. Scale bar, 100 μm. Angiotensin II treatment consisted of 490ng/Kg/day for 14 days, via osmotic mini-pump. Data are presented as mean ± SEM (n =3-4). *P < 0.05 vs PGRN-/-.

## Discussion

Loss of vascular contraction is a critical factor in CVD pathogenesis limiting the normal function of the artery and increasing the risk of organ damage because of impaired blood flow^46^. However, the mechanisms involved in this phenotype are not well understood. In this study, we uncover a novel and major role for PGRN, which consists in maintaining the VSMC contractility via adjusting mitochondrial quality and dependent on lysosome biogenesis, mitophagy flux, and complex I biogenesis pathways.

PGRN loss-of-function mutations cause neuronal ceroid lipofuscinosis and FTD-GRN in a dosage-dependent way^47^. PGRN regulates development, survival, function, and maintenance of mammalian cells including vascular cells (VSMC^48^ and endothelial cells^15^). Recently, Gerrits et al^49^ identified that the neurovascular unit is severely affected in FTD-GRN, while our group described that deficiency in PGRN disturbs endothelial biology and blood pressure regulation ^15^. In VSMC, PGRN exerts antimigratory effects via modulating IL-8 formation in atherosclerotic environment^48^ and its full-length (∼70kDa) acts inhibiting the calcification in calcific aortic valve disease^16^. Therefore, PGRN plays a major role in regulating vascular biology and it seems to be a key protein keeping a contractile VSMC phenotype. Herein, for the first time, we are describing that deficiency in PGRN strikingly affects the contraction of VSMC likely suppressing the amount of contractile proteins and fibrosis and without affecting the vascular structure, at least at 12-16- weeks old mice.

In neuronal^42^ and renal cells^41^ and in *C. elegans^50^* PGRN acts as a regulator of mitochondrial function via preserving mitochondrial dynamics, biogenesis, and mitochondrial recycling. Mitochondrial dysfunction is part of premature ageing and contributes to appearance of inflammation, mitochondria-associated oxidative stress, cell senescence, and apoptosis, which are leading causes of CVD^39,51,52^ including atherosclerosis and aneurysm ^53-55^. From a mechanistic perspective, our observations revealed that the deficiency of PGRN had a multifaceted impact on mitochondrial quality within VSMCs. Notably, VSMCs deficient in PGRN exhibited mitochondrial fragmentation and a decrease in the expression of complex I genes and mass. These alterations subsequently exerted a profound influence on mitochondrial respiration, as evidenced by a reduction in complex I activity, a disrupted NAD+/NADH ratio, and diminished ATP levels. The compromised mitochondrial function culminated in an excessive generation of mtROS. Notably, the reduction in mitochondrial complex I mass, or activity has been recognized as a significant factor contributing to redox imbalance and the development of cardiovascular disease^56^. In our study, we observed that the disruption of PGRN signaling led to an excessive production of mtROS and heightened sensitivity to antimycin A-induced mtROS generation. Significantly, the overexpression of PGRN in vascular smooth muscle cells (VSMC) resulted in the development of high-capacity mitochondria characterized by enhanced mitochondria fusion, increased complex I activity, a higher NAD+/NADH ratio, elevated ATP levels, and improved antioxidant properties (inhibition of mtROS formation). This was followed by VSMC hypercontractility. From a therapeutic standpoint, our findings suggest that the overexpression of PGRN in VSMC could potentially regulate mitochondrial capacity and provide protection against CVD. This protection is likely achieved through precise modulation of mtROS signaling, facilitation of mitochondrial complex I biogenesis, and enhancement of its function. While the specific mechanisms by which PGRN regulates mitochondrial complex I activity and biogenesis were not investigated in this study, previous research in neuroblastoma cell lines has shown that PPARγ activation can rescue mitochondrial function when complex I is inhibited. Therefore, the observed suppression of PPARγ in aortae from both male and female PGRN-deficient mice may indicate impaired formation of genes related to complex I^45^.

Mitochondria are exceptionally dynamic organelles that undergo synchronized cycles of fission and fusion, a phenomenon referred to as mitochondrial dynamics. These processes are essential for maintaining mitochondrial shape, distribution, and size^39,57^. In a positive feedback loop, an excess of reactive oxygen species (ROS) triggers mitochondrial fission^58^, while mitochondria can rapidly fragment, accompanied by an increase in ROS generation^59^. Consequently, within the vasculature, it is plausible that PGRN exerts its antioxidant properties by regulating complex I activity and mitochondrial dynamics, as previously observed in non-vascular cell types^42^.

Deficiency in PGRN regulates the expression of PGC1α in non-vascular tissues^41,60^, in line with this observation, we noted a significant decrease in PGC1α in VSMC deficient for PGRN. Surprisingly, this suppression did not result in changes in mitochondrial quantity. PGC1α is transcriptional coactivator that not only governs mitochondrial biogenesis, but also is linked to antioxidant defense and anti-inflammatory property in endothelial cells and VSMC^43,44,61^. Hence, the reduction in PGC1α may affect inflammatory responses and redox balance rather than mitochondrial biogenesis in PGRN-/- VSMC, and perhaps other mitochondrial regulator might be overcoming the impaired PGC1α, including PGC-1β, which shares similar molecular structure and function with PGC-1α^62^. Further research is necessary to explore the interface between PGRN and both PGC1α and β in cardiovascular biology.

It appears that PGRN exerts its biological effects primarily in its secreted form, as demonstrated previously in endothelial cells^15^. To investigate this further, we sought to restore circulating PGRN levels in PGRN-deficient mice using osmotic mini-pump delivery, as well as in isolated VSMC. Surprisingly, our observations indicated that circulating PGRN did not have a significant impact on mitochondrial function in freshly isolated aortae or in cells treated with rPGRN. These findings strongly suggest that PGRN does not maintain vascular contractility and mitochondrial quality through autocrine or paracrine mechanisms. Instead, it appears that PGRN may function as an intracellular molecule. To delve deeper into this concept, we reconstituted PGRN levels in freshly isolated aortae from PGRN-deficient mice via LV delivery, allowing us to assess mitochondria respiration and vascular contractility. Remarkably, the reintroduction of PGRN in fresh aortae led to the restoration of mitochondrial capacity, particularly through the enhancement of complex I activity, and a significant improvement in vascular contractility. This crucial piece of data strongly indicates that PGRN exerts its regulatory effects on vascular contractility and complex I activity from within the cell, rather than in its circulating form.

Mammalian cells have developed finely tuned mechanisms for quality control to ensure the preservation of functional mitochondria, aligning with the cell’s energy demand^63^. Mitophagy is a selective autophagy process responsible for eliminating accumulated damaged or dysfunctional mitochondria from cells, contributing to the maintenance of mitochondrial homeostasis^63^. In diabetic nephropathy, PGRN is crucial in regulating renal integrity by regulating mitophagy^41^. In VSMC we observed that PGRN deficiency leads to an impaired mitophagy flux (accumulation of p62, LC3I/II, PINK, and PARKIN), while overexpressing PGRN in fresh aortae from mt-Keima mice or in VSMC leads to an accelerated mitophagy process. To explore the potential therapeutic avenue of reinvigorating autophagy in individuals with PGRN deficiency, we treated mice with spermidine, a natural polyamine with autophagic characteristics ^25^and life-span-extending effect in mammalian organisms ^25,53,64^. Spermidine effectively induced autophagy improved the mitochondria function and rescued the vascular function in PGRN deficient mice. These compelling findings suggest that the reinstatement of mitophagy flux holds promise as a pharmacological strategy to mitigate cardiovascular risk in patients with loss-of-function mutations in PGRN. This therapeutic approach offers exciting prospects for addressing cardiovascular concerns in PGRN-deficient individuals.

PGRN is a guardian of lysosome biology ^65^ by regulating lysosome biogenesis in a TFEB dependent manner^66^ and lysosome function. Lack of PGRN triggers accumulation of p62 (marker of damaged lysosome^67^) and LAMP1/2^68,69^. In line with previous findings, PGRN deficient VSMC demonstrated accumulation of p62 and LC3I/II, increased LAMP content, and greater nuclear TFEB levels, although there is more TFEB in the nuclear fraction - an indicative of higher activity - the flux autophagic is interrupted (increased p62 and LC3I/II accumulation), indicating that TFEB is more active, but the formed lysosomes are imperfect. This heightened activation of TFEB may be indicative of an effort to stimulate the formation of suitable lysosomes and subsequently establish a proper autophagy flux, albeit through an inefficient pathway. Interestingly, high amount of PGRN promoted more TFEB translocation into the nuclei and led to a prominent p62 reduction and increased LC3I/II. These observations strongly suggest that PGRN plays a role in triggering TFEB activation within lysosomes, subsequently influencing autophagy flux. Therefore, it becomes evident that complex I activity, biogenesis, and the disturbance of mitophagy are key downstream pathways influenced by PGRN.

Since lack of vascular PGRN pathway is a cardiovascular risk, we challenged mice with Angiotensin II for two distinct reasons 1. to analyze whether ablation of PGRN would aggravate Angiotensin II-induced mitochondria dysfunction and 2. to examine whether Angiotensin II could restore the vascular contractility in PGRN deficient mice. Normally, Angiotensin II treatment induces vascular hypercontractility^70^, upregulates contractile proteins^71^, and triggers mitochondria dysfunction^72^. At least with Angiotensin II treatment, lack of PGRN did not affect mitochondrial function, but it blunted the vascular hypercontractility, thus vascular PGRN pathway is essential to maintaining the vascular tone and induce vascular contraction.

In this study, we propose that the PGRN pathway plays a pivotal role in the regulation of vascular tone through the modulation of mitochondrial quality, utilizing two distinct and sequential mechanisms. PGRN regulates mitochondrial complex I biogenesis and activity and orchestrates mitochondria recycling via maintaining lysosome quality. Subsequently, it preserves vascular bioenergetic pathway and redox balance inhibiting an exacerbated mtROS formation, which conserves the vascular tone. Future research endeavors should delve into the long-term consequences of PGRN deficiency in the context of cardiovascular diseases. Longitudinal studies are needed to assess how PGRN levels correlate with disease severity and progression in human patients. Finally, our study opens new opportunities not only to cardiovascular biology, but also to examine this pathway in neurodegenerative diseases associated with PGRN mutations, which have been recently associated with perturbation of neurovascular compartment^49^, as well.

## ACKNOWLEDGMENTS

The authors are grateful to Dr. Toren Finkel of the Aging Institute at University of Pittsburgh for sharing the mt-Keima mice with our laboratory. The Flow Cytometry Core at the Children’s Hospital of Pittsburgh and Joshua J. Michel for helping us with ROS measurement analysis. This work was funded by NHLBI-R00 (R00HL14013903), American Heart Association Career Development Award (AHA-CDA857268), Vascular Medicine Institute, University of Pittsburgh, the Hemophilia Center of Western Pennsylvania Vitalant, in part by Children’s Hospital of Pittsburgh of the UPMC Health System, and start-up funds from the University of Pittsburgh to Dr Bruder-Nascimento and R01-HD103602 and R01-DK090242 to Dr Eric Goetzman.

## AUTHOR CONTRIBUTIONS

Conceptualization: S.S., E.G., and T.B.N. Methodology: S.S., R.M.C., A.B.N., S.S.B., E.G., and T.B.N. Validation: S.S., R.M.C., A.B.N., and T.B.N. Formal Analysis: S.S., R.M.C., and T.B.N. Investigation: S.S., R.M.C., A.B.N., S.S.B., and T.B.N. Resources: E.G. and T.B.N. Writing – Original Draft: S.S. and T.B.N. Writing – Review & Editing: S.S., E.G., and T.B.N. Visualization: S.S., R.M.C., and T.B.N. Supervision: E.G. and T.B.N. Project Administration: T.B.N. Funding Acquisition: E.G. and T.B.N.

